# Fine-Tuned Deep Transfer Learning Models for Large Screenings of Safer Drugs Targeting Class A GPCRs

**DOI:** 10.1101/2024.12.07.627102

**Authors:** Davide Provasi, Marta Filizola

## Abstract

G protein-coupled receptors (GPCRs) remain a focal point of research due to their critical roles in cell signaling and their prominence as drug targets. However, directly linking drug efficacy to receptor-mediated activation of specific intracellular transducers and the resulting physiological outcomes remains challenging. It is unclear whether the enhanced therapeutic window of certain drugs — defined as the dose range that provides effective therapy with minimal side effects — stems from their low intrinsic efficacy across all signaling pathways or ligand bias, wherein specific transducer subtypes are preferentially activated in a given cellular system compared to a reference ligand. Accurately predicting safer compounds, whether through low intrinsic efficacy or ligand bias, would greatly advance drug development. While AI models hold promise for such predictions, the development of deep learning models capable of reliably forecasting GPCR ligands with defined bioactivities remains challenging, largely due to the limited availability of high-quality data. To address this, we pre-trained a model on receptor sequences and ligand datasets across all class A GPCRs, and then refined it to predict low-efficacy compounds or biased agonists for individual class A GPCRs. This was achieved using transfer learning and a neural network incorporating natural language processing of target sequences and receptor mutation effects on signaling. These two fine-tuned models—one for low-efficacy agonists and one for biased agonists—are available on demand for each class A GPCR and enable virtual screening of large chemical libraries, thereby facilitating the discovery of compounds with potentially improved safety profiles.

## INTRODUCTION

G Protein Coupled Receptors (GPCRs) represent one of the largest families of cell–surface signaling proteins, with approximately 800 members encoded in the human genome (*1*). They are categorized into five families, with the rhodopsin-like GPCR family (class A) being the largest, comprising 289 non-olfactory members across 65 subfamilies, according to GPCRdb (*2*). By responding to a variety of physical and chemical stimuli, GPCRs regulate the function of nearly every organ in the human body, making them critical drug targets for more than one-third of all clinical drugs.

A thorough understanding of the pharmacology of drugs targeting GPCRs requires insight into their capacity to activate distinct signal transduction pathways, as this can lead to diverse physiological outcomes (*3, 4*). Notably, when a ligand preferentially induces receptor activation of one transducer pathway over others compared to a reference ligand within a particular cellular environment ― a concept known as “ligand bias” (initially observed with G protein over β-arrestin pathways) ― it intersects with inherent cellular “system biases.” This combined phenomenon, termed “functional selectivity” or “biased agonism” (*3–9*), has been linked to improved therapeutic outcomes for certain drugs, especially those targeting adrenergic or opioid receptors (see, for example, refs. (*10, 11*)). This has spurred considerable interest in developing GPCR-targeting drugs that elicit functionally selective responses, with the potential to activate therapeutic pathways while minimizing adverse effects.

Recent studies suggest that low intrinsic efficacy — the ability of an agonist to produce a weaker cellular response when engaging its target GPCR compared to a full agonist — rather than biased agonism, which accounts for both the intrinsic efficacy and potency of an agonist in an assay, may contribute to the improved therapeutic windows observed for certain GPCR agonists (*12, 13*). Given that context-dependent differences in cellular systems cannot be ruled out, and a reanalysis of the data questioned this conclusion (*14*), the debate continues on whether the improved therapeutic profiles of certain drugs arise from their low intrinsic efficacy in all signaling pathways or rather from their biased agonism, which may extend beyond the preferential activation of G proteins over arrestins and include different G protein subtypes (*15*).

Recent advancements in AI and machine learning (ML)-based tools, particularly deep learning models, offer promising avenues for identifying ligands with specific bioactivity profiles and physiological outcomes. By rapidly recognizing patterns in vast datasets with high precision (*16*), these tools are significantly impacting drug discovery (*17, 18*), accelerating and improving the process (*19, 20*) through enhanced predictions of GPCR structures, ligand-GPCR interactions, clinical responses, and novel GPCR ligands (e.g., see (*21–24*)). However, the limited size of training datasets remains a challenge for making accurate predictions for certain GPCRs. Transfer learning—an ML technique (*25*) that involves pre-training models on a task for which large datasets are available and then fine-tuning them on tasks with limited data — has proven effective in achieving efficient learning and optimal performance, especially when training on combinations of ligand-based and structure-based descriptors. As we recently demonstrated in the case of opioid receptors, this strategy effectively reduces misclassification and improves the prediction of bioactive opioid ligands at each receptor subtype using dense neural network (DNN) (*26*) or graph convolutional network architectures (*27*).

In this study, we used transfer learning and a deep learning framework that integrates DNN classifiers with the transformer-based ProteinBERT architecture (*28*) to pre-train a model on receptor sequences and datasets of active ligands across all class A GPCRs. We then fine-tuned this model to predict low-efficacy compounds or biased agonists for each class A GPCR with precision. These models, which demonstrate relatively high performance for certain class A GPCR subfamilies, are made available to the scientific community upon request as they hold significant potential for identifying safer compounds through *in silico* screening of large chemical databases—a task too computationally demanding for classical docking methods. These tools will help increase the number of GPCR ligands with characteristics of low-efficacy compounds or biased agonists with potentially improved therapeutic windows, thereby advancing both our understanding of GPCR function and the development of new, safer drugs.

## METHODS

### Deep Learning Framework

We developed a deep learning framework to enhance ligand activity prediction for class A GPCRs. This framework integrates: (i) embeddings of GPCR sequences and functions generated using the transformer-based ProteinBERT model (depicted in grey and brown in Figure 1), and (ii) embeddings of ligands generated through a DNN trained on ligand structural fingerprints (depicted in blue in Figure 1). These receptor and ligand embeddings are used as inputs to a DNN (illustrated in pink in Figure 1) to predict ligand-receptor activities.

**Figure 1.**
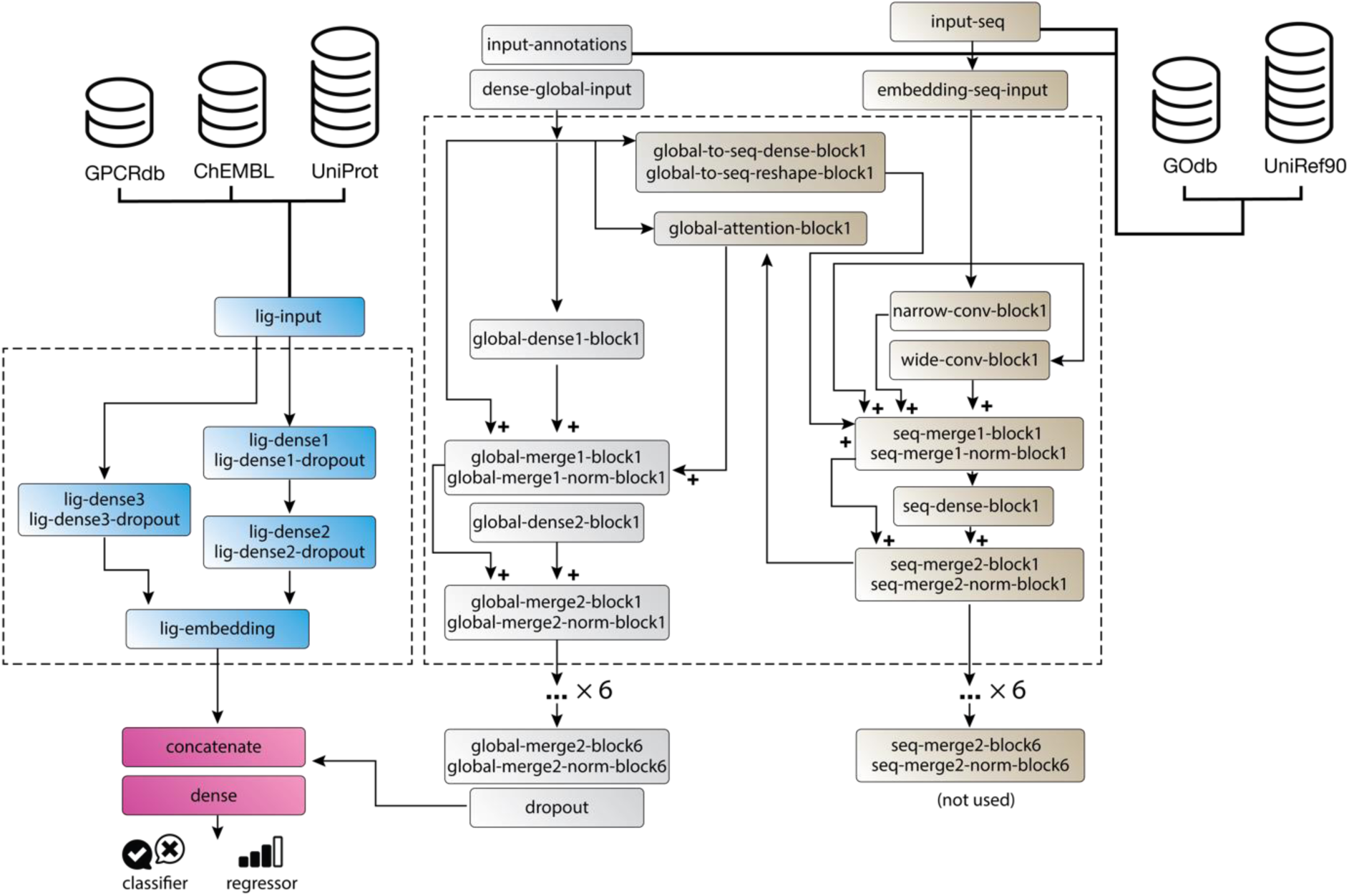
Deep Learning Framework for Predicting Ligand-Receptor Activities. Dense layers generate a ligand embedding (blue) from structural fingerprints, while the ProteinBERT transformer (brown and grey) produces a global embedding of the target GPCR based on its functional annotation and sequence. The combined ligand and target GPCR representations are fed into a DNN model, which outputs a classification label (e.g., active vs. inactive or full vs. partial agonist) or predicts a (log) activity ratio.

Specifically, we trained a potency-based classifier to predict whether a given ligand is active at a specific GPCR, and an efficacy-based classifier to determine whether an active ligand functions as a full or partial agonist. Additionally, we developed an activity regressor to estimate the average “activity ratio” (E_max_/EC50)^*i*^ (*29*) for G protein and arrestin signaling endpoints *i*. Using the activity regressor’s prediction for different signaling pathways *i* and *j*, it is possible to estimate ligand bias in terms of the relative intrinsic activity difference between a ligand *a* and a reference ligand *b* (*14*), expressed as ΔΔlog(*E*_*max*_/*EC*50)_a,b_^i,j^. Throughout the paper log indicates the logarithm in base 10.

For protein embeddings, we used ProteinBERT version 1.0.1, a large language model with six transformer blocks that capture both local (sequence-specific) and global (feature-based) protein representations derived from the UniRef90 and Gene Ontology (GO) databases, respectively, using two interconnected processing tracks. Each transformer block links the global protein representation, *x* (grey in Figure 1), with the local sequence representation, *s*_*i*_ (brown in Figure 1), via fully connected layers. The local track refines the protein sequence embedding using narrow and wide convolution blocks with strides of 1 and kernel sizes of 9 and 41, respectively, followed by dense layers (see Figure 1 for details).

Connections between the local and global representations are established through *global* attention layers. Specifically, embeddings *s*_*i*_ for each residue *i* in the target protein sequence are converted to a local key, *k*_*i*_ = *σ*(*W*_*k*_*s*_*i*_), and values, *v*_*i*_ = *σ*(*W*_*v*_*s*_*i*_), while the *global* protein embedding *x* is transformed into a global query, *q* = *σ*(*W*_*q*_*x*), where *W*_*k*_, *W*_*v*_, and *W*_*q*_ are learned transformation matrices. The local key, global query, and value vectors are then used to compute an attention score, *y* = 〈*v*_*i*_,softmax 〈*q*, *k*_*i*_〉〉. This score highlights how each sequence position aligns with global properties, elucidating which residues contribute crucial information for predicting the receptor’s global features.

For ligand embedding, ligands were represented as *Simplified Molecular-Input Line-Entry System* (SMILES) strings, which were converted to their canonical form using RDKit’s `CanonSmiles` function (*30*). The molecular structures were then fingerprinted using Morgan’s extended-connectivity fingerprinting approach (ECFP4) (*31, 32*) with a radius of 2 to obtain 1024- bit descriptors generated via RDKit’s `GetMorganGenerator` and `GetFingerprint` functions. These structural descriptors were used as input for generating ligand embeddings through two parallel functions: one comprising two dense layers with ReLU activation and dropout, and the other consisting of a single ReLU layer with dropout (illustrated in blue in Figure 1). The resulting ligand embeddings were then combined with the protein embeddings and processed through a final dense layer with a Softmax activation function to compute class probabilities for ligand activity in the pre-training phase of the potency-based classifier and for partial or full agonism in the fine-tuning phase of the efficacy-based classifier. For ligand bias prediction, the same architecture was used with a linear activation function to predict standardized values of the logarithmic activity ratio, log(E_max_/EC50)^*i*^. To enable robust inference, the regressor was equipped to estimate heteroscedastic aleatoric uncertainty, capturing varying levels of noise inherent in the training observations. Thus, instead of predicting a single value for log(E_max_/EC50), the model employed two parallel final dense layers to estimate the parameters of a normal distribution of the log(E_max_/EC50) ∼ *N*(*μ*, *σ*) (*33*).

### Datasets and Training

A transfer learning approach was employed to pre-train a model capable of distinguishing active from inactive ligands across all class A GPCRs using receptor sequence and ligand bioactivity data, and then fine-tuning it to discriminate between partial and full agonists, as well as to predict activity ratio for different pathways, enabling the identification of biased and balanced agonists for each individual class A GPCR. Following the original ProteinBERT methodology (*28*), a language model was pre-trained on 106 million protein sequences from UniRef90, a curated dataset of UniProtKB protein clusters with at least 90% sequence identity (*28*). The 8,943 most common GO annotations associated with UniRef90 proteins were utilized as global protein features for self-supervised pre-training. These annotations covered: (i) *Molecular Function*, describing molecular-level activities performed by the target (e.g., catalytic or transcription regulator activity); (ii) *Cellular Component*, indicating the cellular location of these activities, such as cellular structures and protein complexes (e.g., mitochondrion or plasma membrane); and (iii) *Biological Process*, representing broader biological programs that integrate multiple molecular functions (e.g., DNA repair or signal transduction). Tokens representing the twenty natural amino acids, along with “U” for selenocysteine and “X” for unknown residues, as well as special tokens for the “start” an “end” of the sequence, a “pad” token, and an “other” token, were used to represent the local features of the protein sequence. During self-supervised pre-training, the model was presented with corrupted inputs (protein sequences and GO annotations) and tasked with reconstructing the original data. The training in this phase employed a loss function that combined the categorical cross-entropy across protein sequences with binary cross-entropy for the GO annotations, allowing the language model to learn protein representations that effectively capture the relationships between protein sequence and function.

#### Training the Model for Potency-Based Activity Classification

After pre-training on the 106 million protein sequences from UniRef90, the full model architecture, including the ligand classifier, was trained to distinguish active from inactive class A GPCR ligands. All bioactivity data for class A GPCRs, excluding olfactory GPCRs, were sourced from the ChEMBL 34 database (*34*).

Bioactivities with *“*standard type” annotations of “IC50”, “EC50”, or “AC50” and a “standard value” threshold of 1 μM were used to classify ligands as either active or inactive. The resulting potency-based activity dataset included 168,822 inactive and 214,816 active ligand data points, totaling 383,638 GPCR-ligand pairs. A comparison of this dataset’s size with other datasets used in this study is provided in supplementary Table S1. This dataset represented 180,340 unique ligands (ChEMBL molecule IDs) and 615 unique targets (Target ChEMBL ID) from 28 different organisms, with a predominant representation of *Homo Sapiens* (80%) and *Rattus norvegicus* (11%). Wild-type receptor sequences were sourced from UniProt, cross-referenced with target gene labels in ChEMBL, and filtered to exclude sequences exceeding 500 residues to enhance training efficiency (*28*). Additionally, information on receptor mutations affecting GPCR signaling was incorporated from GPCRdb (*35*) to further enrich the dataset. Initially, the model was trained for 10 epochs on the potency-based activity dataset with the parameters of the ProteinBERT layers kept fixed. A second phase of fine-tuning was then performed for 10 epochs, during which all weights in the model were adjusted. During this phase, the learning rate was reduced whenever validation loss plateaued, using the Keras `ReduceLROnPlateau` function with a patience of 1 epoch and a reduction factor of 0.25. To prevent overfitting, early stopping was applied using the Keras `EarlyStopping` function with a patience of 2 epochs.

*Fine-Tuning the Model for Low-Efficacy Classification.* To classify ligands based on efficacy, we selected all ChEMBL bioactivities with an *Emax* “standard type” and “standard units” expressed as a percentage, focusing on ligands previously identified as actives in the potency-based classification dataset. Ligands were categorized as “full-agonists” if the reported *Emax* exceeded 70% and as “low-efficacy/partial-agonists” otherwise. To address the possible misclassification of ligands with a reported Emax of 100% as full agonists rather than partial agonists due to signal amplification in certain assays caused by significant receptor reserve, we applied the Black and Leff operational model (*36*). According to this model, EC50 = *K*_*A*_/(*τ* + 1), where *τ* is a measure of the intrinsic efficacy and *K*_*A*_ is the equilibrium constant for ligand binding to the receptor. For ligands with low intrinsic efficacy, EC50 ≈ *K*_*A*_ (*37*). Based on this principle, ligands with both potency values (e.g., standard types of EC50, IC50, AC50) and binding affinity values (e.g., Ki, Kd) available in ChEMBL were analyzed. Ligands classified as full agonists with an *Emax* = 100% but showing a potency-to-affinity ratio between 90% and 100% were excluded, as these values suggested that, in non-amplified assays, the ligands might act as partial agonists. The final efficacy-based dataset (Table S1) included 6,278 full agonists and 4,193 partial agonists, totaling 10,471 ligand-GPCR pairs. These pairs comprised 7,749 unique ligands and 198 unique targets from 17 organisms, with *Homo sapiens* (85%) and *Rattus norvegicus* (7.6%) being the most represented. A breakdown of the dataset by class A GPCR subfamilies with the largest number of data points is provided in Table S2. The model underwent two training phases, beginning with the model pre-trained on the potency-based classifier. The first phase consisted of 10 epochs, during which the weights of the protein and ligand embeddings were kept fixed. This was followed by a fine-tuning phase of 10 epochs, during which all layers were optimized.

#### Fine-Tuning the Model for Ligand Bias Assignment

Data on biased agonists were sourced from the online Biased Signaling Atlas (*38*). Only data points reporting both EC50 (or pEC50) and Emax values were retained for analysis. Data points annotated with effector family Gq/11, Gi/o, G12/13, Gs, or “G protein” were grouped into the “G protein” class, while those annotated as ERK or arrestin were assigned to the “arrestin” class. Potency values were converted to uniform units, and the log activity ratio was calculated using molar EC50 values.

The final dataset comprised 6,085 data points, including 1,030 unique ligands and 64 unique receptors. This total included 2,304 datapoints (864 unique ligands and 56 unique targets) for arrestin pathway effectors and 3,781 datapoints (992 unique ligands and 62 unique receptors) for G protein pathways. The range of the logarithmic activity ratio, log(E_max_/EC50), was 2.22-11.3 for arrestin pathways and 1.14-11.2 for G protein pathways.

#### Validation Sets and Performance Metrics

All training was conducted with at least five independent random splits of the data into training (80%) and validation (20%) sets, ensuring both active and inactive classes were randomly split. Additionally, model performance was assessed on validation datasets obtained through two stratification methods: (1) stratifying by ligands (“ligand split”) so that 20% of the ligands were exclusively assigned to the validation set and excluded from training, and (2) stratifying by targets (“target split”) so that 20% of the targets were assigned exclusively to the validation set. For the potency-based and efficacy-based classification tasks, we report the following performance metrics: (i) Precision (also called positive predictive value): the fraction of correctly predicted positive instances (i.e., true actives for the potency-based classifier or true low-efficacy instances for the efficacy-based classifier) among the retrieved instances (i.e., predicted actives for the potency-based classifier or predicted low-efficacy for the efficacy-based classifier); (ii) Recall (also known as sensitivity or true positive rate): the fraction of actual positive instances that the model correctly identifies; (iii) F1 score: the harmonic mean of precision and recall; and (iv) AUCROC: the Area Under the Receiver Operating Characteristic (ROC) curve, which represents the trade-off between recall and the false positive rate as the threshold varies from 0 to 1. Performance metrics for pre-training on the potency-based classification task calculated on the random, ligand, and target validation splits are reported in Table S3 whereas Table S4 presents the corresponding metrics for the fine-tuned efficacy classification task, or the classifier trained directly on the efficacy dataset. Table S5 lists F1 scores for the potency-based and efficacy-based classifiers on random validation sets by target family.

For the activity ratio regression models, we report the following performance metrics: (i) The root mean square error 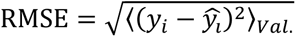; (ii) The mean absolute error MAE = 〈|*y*_*i*_ − *y*^_*i*_|〉_*val*._; (iii) The Pearson and Spearman correlation coefficients, which measure linear and rank-order correlation, respectively; and (iv) The concordance index (CI), which is a metric commonly used to evaluate the model performance of regressors for drug predictions. CI measures the fraction of compounds pairs (*i*, *j*) for which the predicted order of the variable agrees with the true order. Specifically, it calculates the fraction of ligand pairs *i*, *j* for which the relative activity index log(*RA*^*i*^/*RA*^*j*^) > 0 is correctly predicted as log(*RA*^^*i*^/*RA*^^*j*^) > 0. Table S6 reports these performance metrics for the arrestin and G protein pathways.

## RESULTS AND DISCUSSION

### Potency-Based Classifiers Accurately Predict Active Ligands for Class A GPCR

To address the challenge of limited high-quality data for building reliable deep learning models to predict GPCR ligands with specific bioactivities, we implemented a transfer learning strategy. We first pre-trained a model on a well-resourced task: distinguishing active from inactive ligands using receptor sequences and ligand activity data across all class A GPCRs. Given the variability in ChEMBL data —where noise arises from inconsistent measurements for the same compound across multiple assays (*39*), we adopted a classification approach that is relatively more robust than precise potency predictions, and used a potency cutoff of 1 μM to label ligands as active or inactive at each given GPCR target (see details in Methods section). This approach leveraged relatively abundant potency data, encompassing ∼0.38 million ligand-GPCR pairs, ∼180,000 unique ligands (ChEMBL molecule IDs), and 615 unique non-olfactory class A GPCR targets (see Table S1). The model demonstrated strong performance in classifying active vs inactive ligand-GPCR pairs after pre-training, achieving an average precision of 0.80 (0.80, 0.80) and recall of 0.88 (0.87, 0.88) on random splits (see Figure 2 and Table S3). Performance was also evaluated on more challenging scenarios: on ligand splits, the model achieved precision and recall of 0.76 (0.76, 0.76) and 0.89 (0.88, 0.89), respectively; on target splits, it achieved 0.71 (0.71, 0.73) precision and 0.80 (0.79, 0.82) recall. These results highlight the model’s capability to generalize and predict signaling activity for ligands and targets that are not included in the training set.

**Figure 2.**
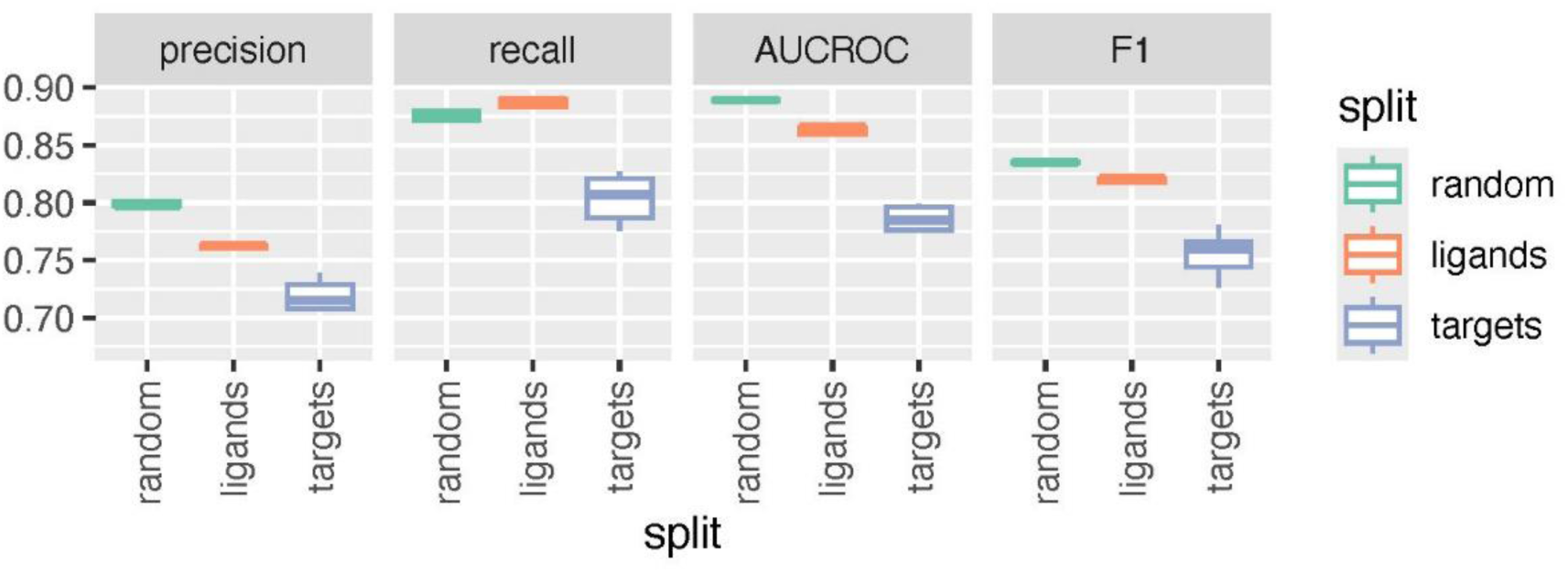
Performance of the potency-based classifier on random, ligand, and target 80%/20% splits, in green, orange, and violet, respectively. The box plots are derived from 5 independent splits and calculated on the validation set.

### Transfer Learning and Protein Language Models Enhance the Prediction of Partial Agonism At Class A GPCR

Building on the potency-based classifier, we fine-tuned this model for tasks with more limited data, particularly those aimed at distinguishing between low-efficacy/partial agonists and full agonists for specific GPCRs. Following a similar approach to the potency-based classification, ligands in the training set were labeled based on a cutoff of 70% for Emax values (see Methods section for details). It is important to note that the Emax values used are: (a) measured via assays probing different endpoints of GPCR function, such as G protein activation or β-arrestin signaling; (b) reported as percentages relative to potentially different full agonists; and (iii) reflect assay- and cell-specific factors, such as target density and coupling efficiency, which vary across experimental conditions. As a result, the predicted efficacy classifications represent average properties of ligand-GPCR pairs rather than true intrinsic ligand efficacy.

The efficacy classifier was trained using a dataset of 10,471 ligand-GPCR pairs, including 7,749 unique ligands (ChEMBL molecule IDs) and 198 unique non-olfactory class A GPCR targets (see Table S1). Starting from the pre-trained potency classifier, we trained the final dense layers of the classifier with constrained weights for the ligand and protein embedding layers for 10 epochs, followed by an additional 10 epochs of fully unconstrained optimization. The efficacy-based classifier’s performance is summarized in Figure 2 (green boxes) and Table S4. For random splits, precision and recall reached 0.70 (0.70, 0.70) and 0.75 (0.72, 0.78), respectively. Performance on ligand splits was comparable, with a precision of 0.71 (0.71, 0.72) and recall of 0.74 (0.73, 0.75). However, for cold target splits, performance decreased significantly, with a precision of 0.57 (0.53, 0.61) and a recall of 0.52 (0.41, 0.61), indicating that the limited data negatively impact ligand classification at receptors not included in the training dataset.

To evaluate the effectiveness of transfer learning in improving model performance, we compared results from models using transfer learning to those from models trained directly on the efficacy-based training set, bypassing pre-training on the potency-based classification task. The results, illustrated in orange in Fig. 3 and summarized in Table S4, confirm the benefits of transfer learning. Precision and recall metrics showed noticeable degradation when pre-training was omitted, across both random and ligand-based data splits. For the random split, precision dropped from 0.70 to 0.66 (0.65, 0.67), and recall decreased from 0.75 to 0.71 (0.69, 0.72). Similarly, on the ligand split, precision declined from 0.71 to 0.66 (0.64, 0.68), and recall fell from 0.74 to 0.69 (0.65, 0.73). Notably, the impact of pre-training was less pronounced on the target cold split, where precision and recall remained relatively stable, decreasing only slightly from 0.57 to 0.56 (0.54, 0.58) and from 0.52 to 0.49 (0.46, 0.48), respectively.

**Figure 3.**
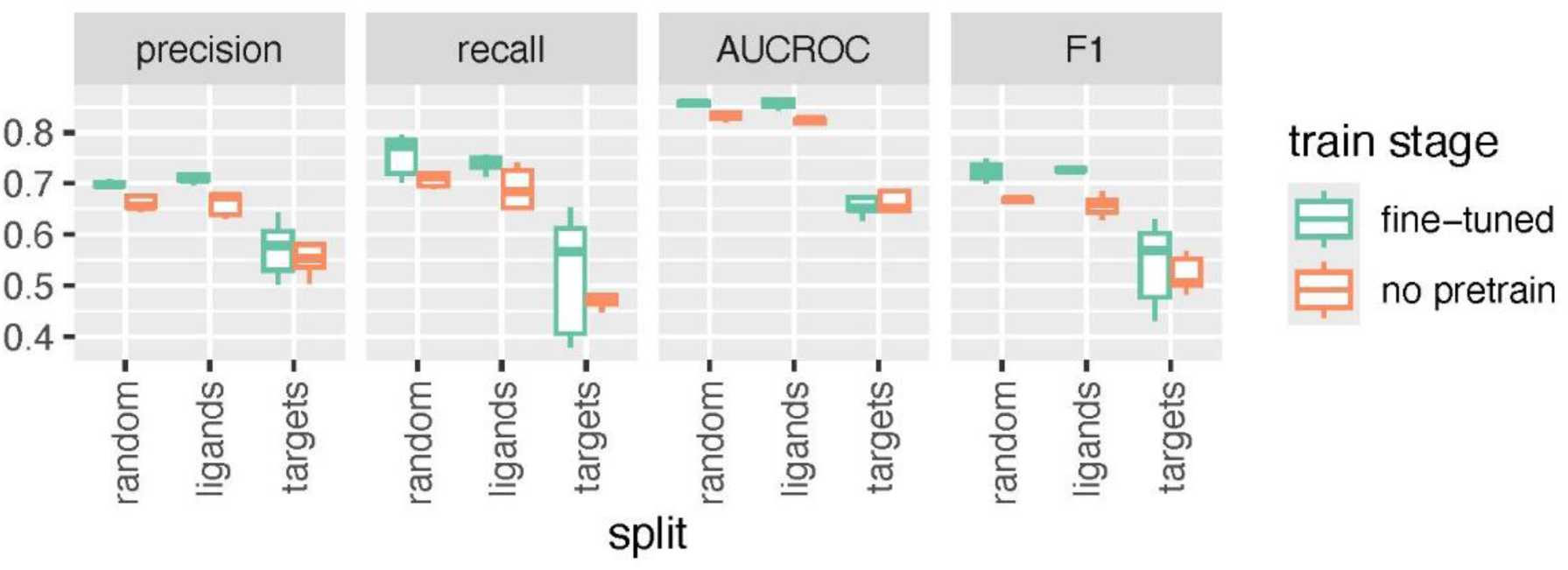
Performance of the efficacy-based classifiers—random, ligand, and target 80%/20% splits—for models fine-tuned after pre-training on the potency-based classification task (green) and those trained directly on the efficacy-based classification task (orange). The boxes represent results from 5 independent splits, calculated on the validation set.

### Predictions of Activity and Partial Agonism Vary in Accuracy Across GPCR Subfamilies

While the aggregated performance metrics presented in the previous sections provide an overview of the average accuracy of potency-based and efficacy-based classifiers, their predictive performance varies considerably across receptor subfamilies. Figure 4 highlights the models’ performance for selected class A GPCRs among the 289 receptors analyzed in this study. Specifically, the figure illustrates the performance variability observed among class A GPCR subfamilies included in at least 4 out of 5 random splits of the validation sets, as measured by F1 scores. These scores, which combine recall and precision, are averaged across all targets within each class A GPCR subfamily. The corresponding values are detailed in Table S5.

**Figure 4.**
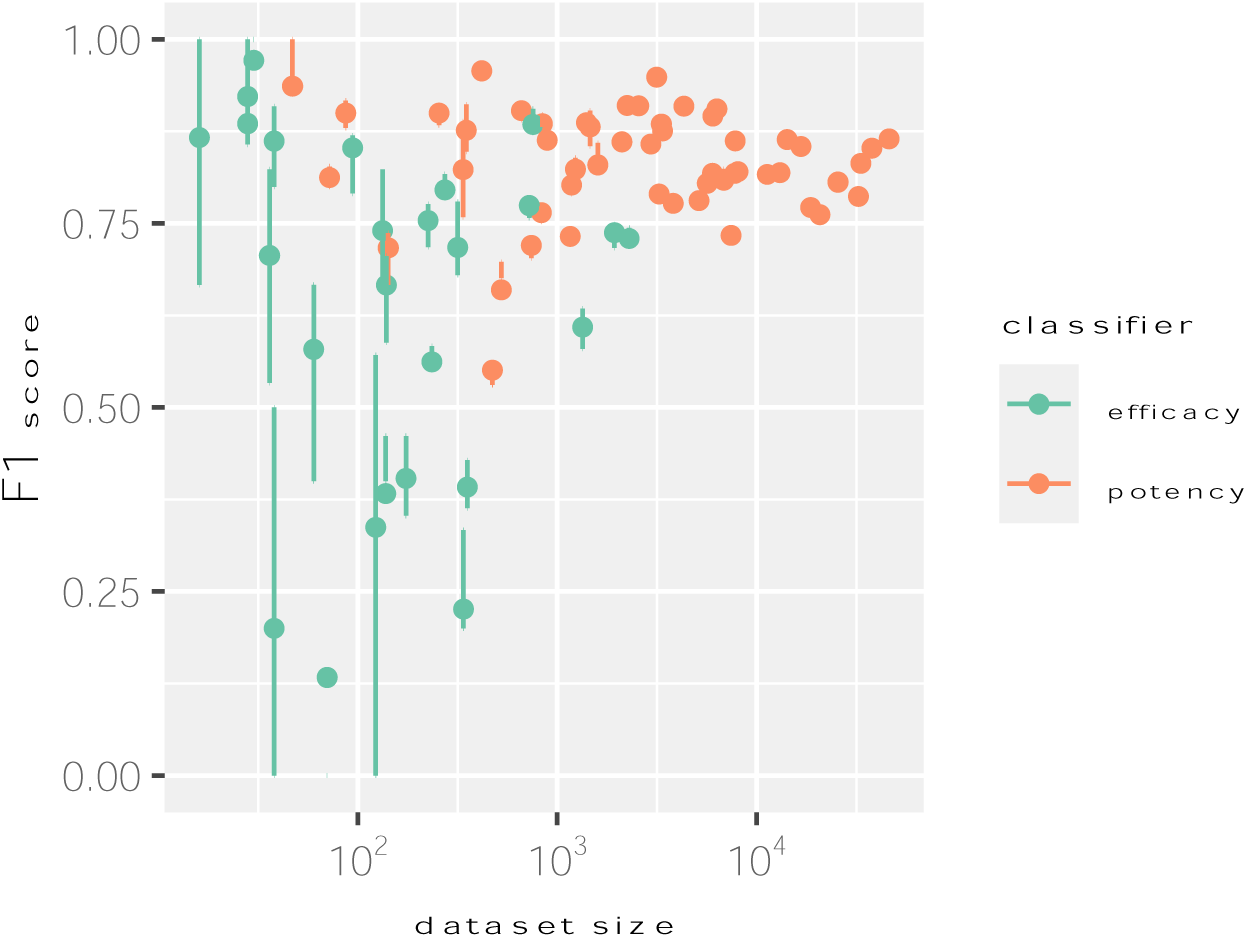
Performance of the classifier models as a function of the size of the training dataset. The average F1 scores over the validation sets of independent splits, along with the 25%-75% intervals, are shown for the potency-based and efficacy-based classifiers in orange and green, respectively.

Notably, the average performance is not directly correlated with the size of the training dataset. To better understand the models’ performance in different contexts, we focus our discussion here on GPCR subfamilies for which biased ligands have been proposed as useful clinical or research tools (*40*). As expected, models’ performance for most receptor subfamilies with relatively large training datasets (i.e., more than 500 data points) is high, achieving F1 scores above 0.73 for the potency-based and efficacy-based classifiers. For instance, the F1 scores for the efficiency-based classifier are 0.74 (0.72, 0.75) for 5-HT receptors, 0.73 (0.73, 0.74) for opioid receptors, and 0.77 (0.76, 0.77) for dopamine receptors. An exception are models obtained for the cannabinoid receptor subfamily, which has a dataset for the efficacy-based classifier comparable to the aforementioned subfamilies but perform significantly worse, with an F1 score of 0.61 (0.58, 0.63). Models obtained for receptor subfamilies with intermediate-sized efficiency-based datasets (i.e., 100-500 datapoints), such as the adenosine receptors, adrenoreceptors, and histamine receptors also show good performance, with F1 scores of 0.80 (0.81, 0.82), 0.72 (0.68, 0.78), and 0.74 (0.67, 0.82), respectively. Notably, performance can also be relatively good for efficacy-based classifiers obtained for receptor subfamilies with smaller datasets (i.e., less than 100 data points). For instance, efficiency-based classifiers obtained for chemokine receptors, prostanoid receptors, and angiotensin receptors achieve F1 scores of 0.71 (0.53, 0.82), 0.97 (0.97, 1.00), and 0.86 (0.80, 0.91), respectively, whereas the apelin receptor model demonstrates particularly weak performance, with an F1 score of 0.17 (0.00, 0.17).

### Few-Shot Fine-Tuning Enhances Partial Agonism Prediction For Selected Class A GPCRs

Performance metrics were initially assessed using zero-shot predictions, where no additional fine-tuning was applied to the model. Few-shot fine-tuning, a transfer learning technique that leverages a small number of samples for model refinement, was subsequently employed. Specifically, the efficacy model was fine-tuned using 10% of the data for a given class A GPCR subtype, resulting in improved predictive performance for selected targets.

The impact of few-shot training is illustrated in Figure 5 using selected targets as representative examples, highlighting variations in training dataset sizes and performance levels. In the figure, violet represents few-shot performance, while orange indicates zero-shot performance. We first notice that few-shot fine-tuning marginally enhances model performance for targets with large training sets and good performance. For example, in the opioid receptor subfamily, the model’s F1 score for the efficacy-based classifier improved from 0.81 (0.80, 0.82) to 0.85 (0.85, 0.86) for OPRM, from 0.63 (0.61, 0.67) to 0.75 (0.75, 0.75) for OPRK, and from 0.67 (0.65, 0.70) to 0.72 (0.72, 0.73) for OPRD.

**Figure 5.**
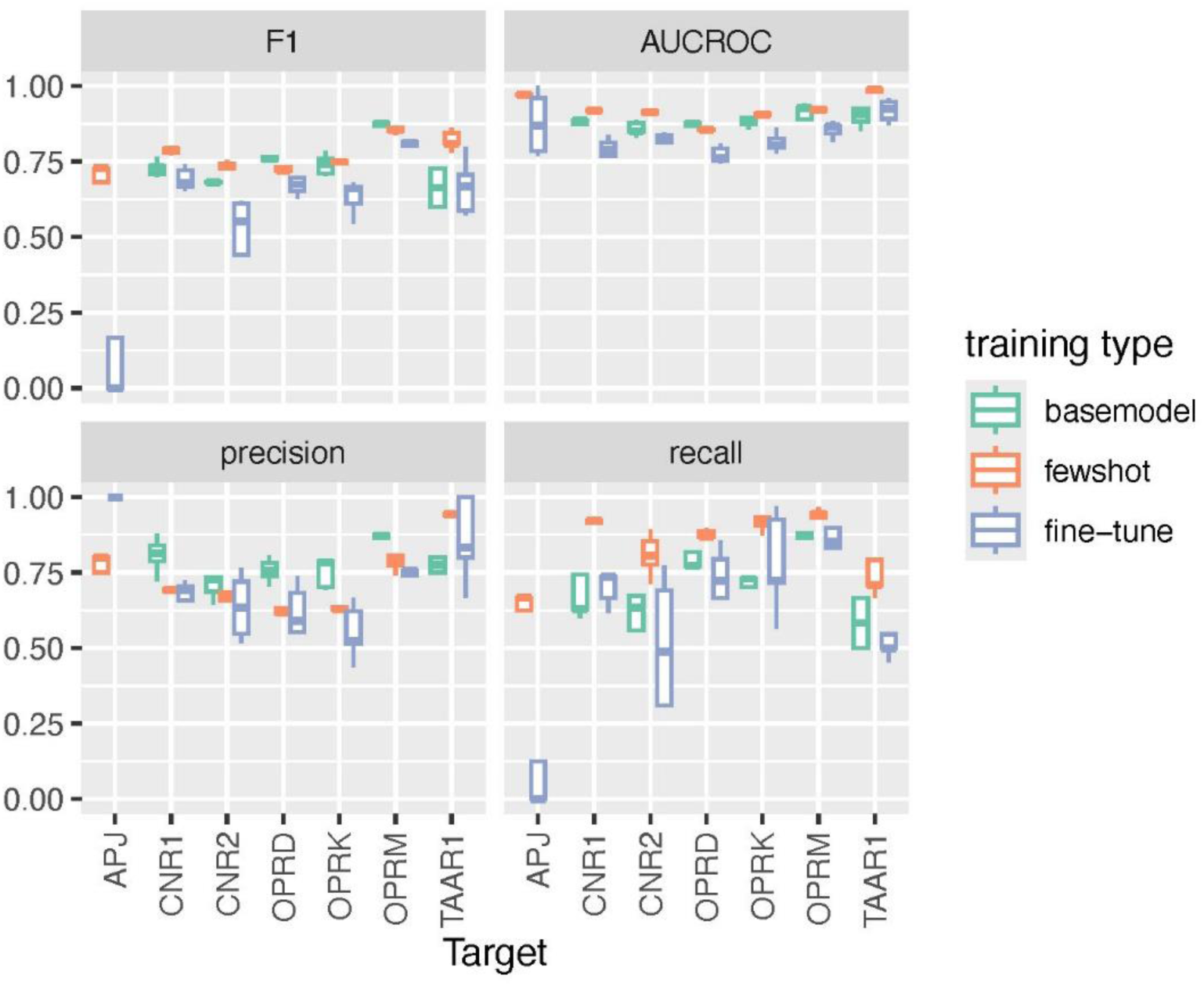
Performance metrics for efficacy-based classifiers for selected targets. The data are presented for three models: the zero-shot application of the protein language model after fine- tuning on the efficacy-based dataset, the few-shot fine-tuned model, and a baseline model trained on each individual receptor without using the language model (shown in violet, orange, and green, respectively). Results are reported for the apelin receptor (APJ), two cannabinoid receptors (CNR1 and CNR2), the three main opioid receptors (OPRM, OPRD, OPRK), and the trace amine receptor (TAAR1). Boxplots are generated using five independent random validation splits.

The few-shot strategy was also effective for targets with large training datasets but suboptimal performance. For example, few-shot training improved the F1 score for the efficacy-based classifier of the cannabinoid CNR1 receptor from 0.69 (0.67, 0.72) to 0.79 (0.78, 0.79), while the F1 score for the efficacy-based classifier of cannabinoid CNR2 increased from 0.53 (0.44, 0.61) to 0.74 (0.73, 0.74). Notably, this strategy is most effective for building predictive models for receptor subfamilies that perform poorly in zero-shot applications, particularly with mid-sized and small training datasets. For instance, on mid-sized datasets (100–500), such as the Trace Amine-Associated Receptor 1 (TAAR1), the F1 score for the efficacy-based classifier improves from 0.67 (0.62, 0.72) to 0.81 (0.80,0.81). Similarly, a brief training on 10% of the small training set of apelin receptor (APJ) improves the F1 score for the efficacy-based classifier on the validation set from 0.17 (0.00, 0.17) to 0.68 (0.68, 0.73).

### Protein Language Models Enhance Efficacy-Based Classifier Performance Compared to Target-Specific Baseline Models

We investigated whether incorporating a protein language model and training on a larger dataset with few-shot fine-tuning could outperform baseline classifiers trained individually for each receptor. As illustrated in Fig. 5, where the performance of the baseline models is shown in green, the results demonstrate improved performance for selected GPCRs when using the protein language model. Not surprisingly, for cases with small datasets, such as APJ, the baseline model fails to predict true negatives, but it is rescued by the protein language model with the few-shot training further enhancing the performance of the zero-shot model, as discussed in the previous section.

For less challenging cases with larger datasets, the baseline models perform comparably to the protein language models with few-shot training. This is exemplified by the three opioid receptors, where the baseline models achieve F1 scores for the efficacy-based classifier of 0.76 (0.76, 0.76), 0.74 (0.71, 0.76), and 0.87 (0.87, 0.88) for OPRD, OPRK, and OPRM, respectively. For other cases such as the cannabinoid receptors CNR1 and CNR2 and the trace amine receptor TAAR1, the protein language model with few-shot fine-tuning consistently outperforms the baseline models, achieving F1 scores for the efficacy-based classifier of 0.73 (0.71, 0.73) and 0.67 (0.68, 0.68) for CNR1 and CNR2, and 0.66 (0.60, 0.73) for TAAR1.

### Prediction of Ligand Activity Ratios (Ligand Bias) Across Class A GPCRs

We next focused on developing models to predict ligand activity ratios, a critical metric for identifying biased ligands. Despite limited data availability, curated datasets provide opportunities for modeling. Similar to the challenges faced in efficacy prediction, variations in assay endpoints—often amplified through distinct cellular signaling pathways—complicate ligand bias prediction. Given these complexities, it is essential to adopt methods that rely on relative activity comparisons rather than directly categorizing ligands as biased or unbiased, as such direct classifications lack meaningful context and render classification-based strategies impractical. To address this, we used Ehlert’s “activity ratio” (sometimes denoted RA) (*29*), which is a useful measure of agonist activity expressed as the ratio of the agonist’s maximal response (Emax) to the concentration required to achieve its half-maximal response (EC50) (*41, 42*).

Using data from the community-curated Biased Signaling Atlas (*38*), we prepared training datasets for deep regression models aimed at predicting the logarithmic activity ratio log (*RA*_*i*_) across assays with varying primary effectors *i*. The effectors were grouped in a “G protein” class, which included G protein families such as Gi/o (2340 data points), Gs (879 data points), G protein (603 data points), Gq/11 (526 data points), and G12/13 (44 data points), and an “Arrestin” class, which included non-G protein effectors such as arrestin (2239 data points) and ERK (461 data points). We began with a model pre-trained to classify ligands based on their potency and trained a regression layer to predict log (*RA*_*i*_). In Figure 6, we present the predictions on the validation set for 5 random cross-validation splits. The accuracy of the predictions are quantified in Table S6 and Figure 7, where we report standard regression metrics.

**Figure 6.**
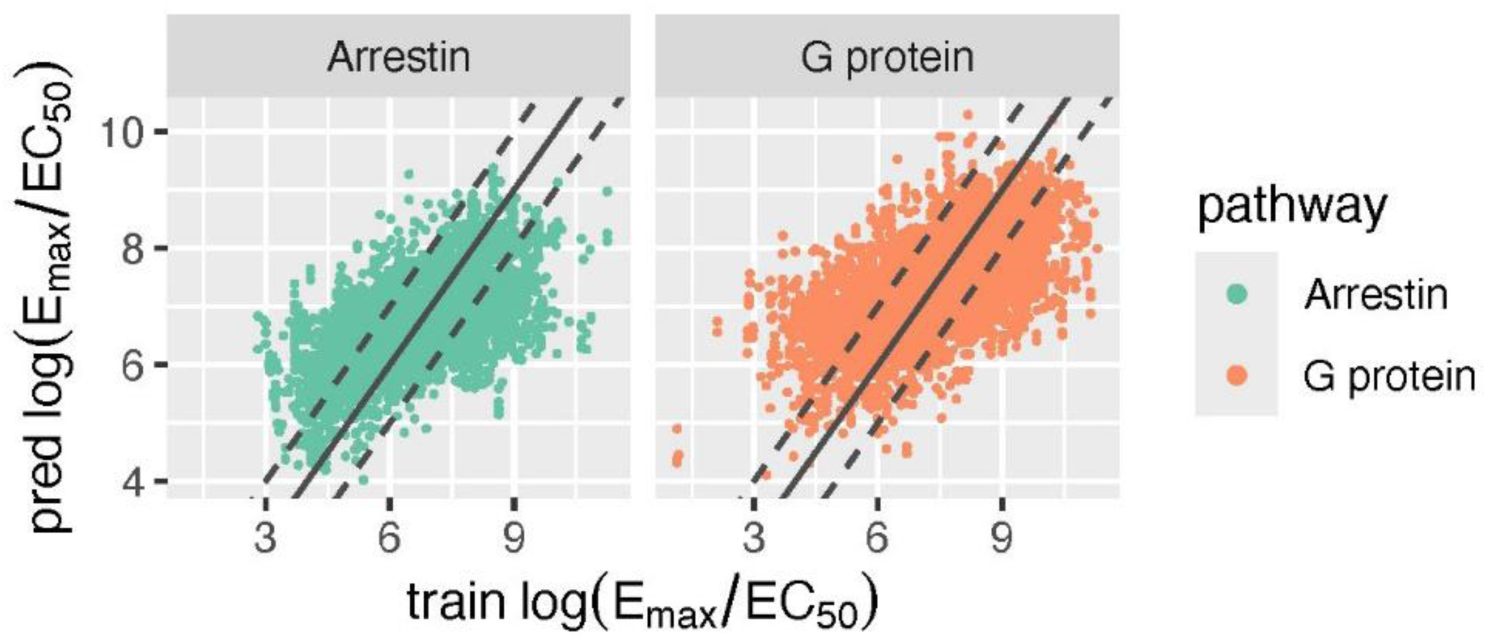
Predicted activity ratio values compared with their corresponding experimental dataset values. Data from the validation sets of 5 random cross-validation splits are presented. Predicted activity ratios for the arrestin and G protein pathways are shown in green and orange, respectively. The solid line represents perfect predictions, while the dashed lines indicate a difference of one unit in the predicted log activity ratio.

**Figure 7.**
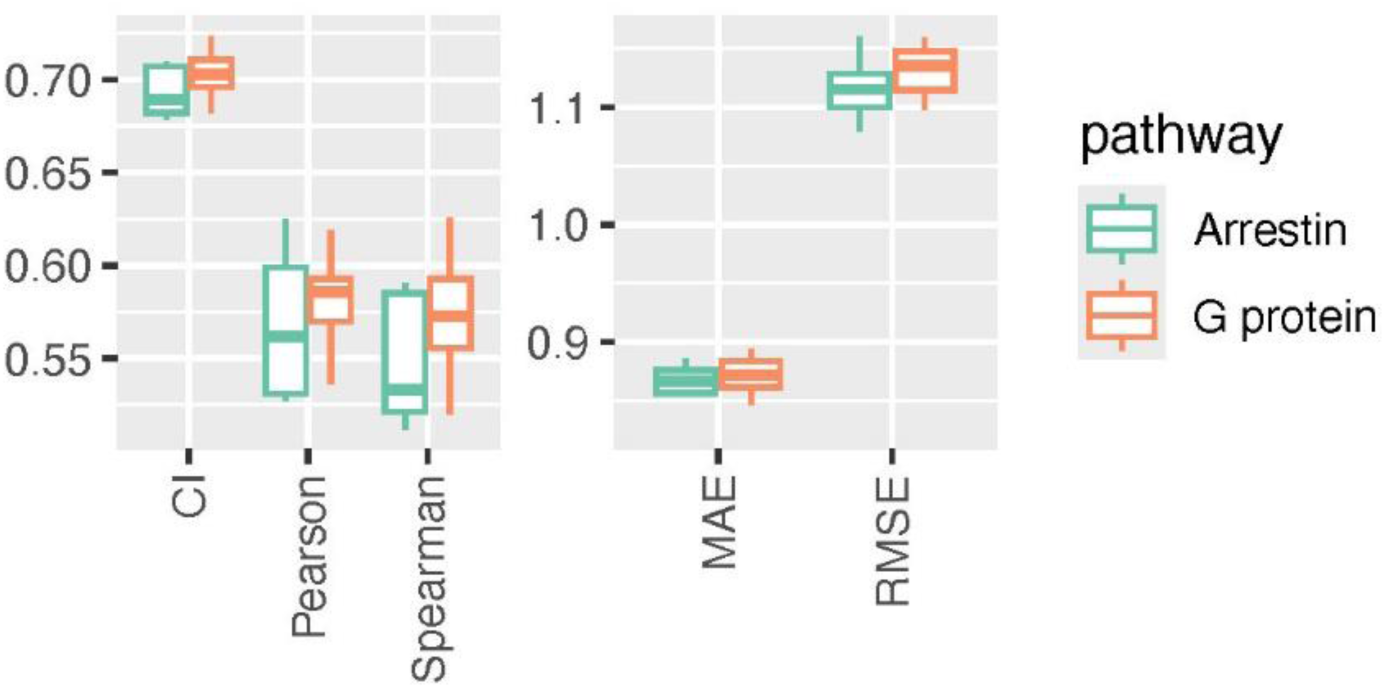
Regression metrics for predicting activity ratio values. Data from the validation sets of 5 random cross-validation splits are reported. Predictions of activity ratio values in the arrestin and G protein pathways are represented in green and orange, respectively.

The regression model for activity ratio predictions in the two signaling pathways performs moderately well, with a MAE of 0.86 (0.86,0.88) for the arrestin pathway and 0.87 (0.86,0.88) for the G protein pathway. The Pearson correlation coefficients are 0.57 (0.53,0.60) and 0.58 (0.57,0.59) for predictions in the arrestin and G protein pathways, respectively. Values of other metrics are reported in Table S6.

The absolute values of log (*RA*_*i*_) cannot be used to assess the bias profile of ligands directly. Instead, after calculating the relative values Δlog(*E*_*max*_/*EC*50)^*i*,*j*^= log(*E*_*max*_/*EC*50)^*j*^ − log(*E*_*max*_/*EC*50)^*i*^, a reference ligand *b* must be selected and the prediction for the two different signaling pathways *i* and *j* must be combined to obtain the relative intrinsic activity difference between a test ligand *a* and the reference ligand b (*14*): 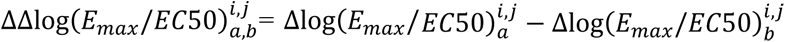. Positive values of ΔΔlog*RA* indicate a bias towards pathway *j*, while negative ones indicate a bias towards *i*. Therefore, for a given reference ligand *b*, a ligand bias classification can be created by assigning each test ligand *a* to a class based on the value of ΔΔlog*RA*.

Here, we evaluate the performance of such classifiers on two targets, OPRM and D2DR, for which biased agonism is an active research focus. For OPRM, we selected DAMGO as the reference ligand, while for D2DR we selected dopamine as the reference ligand. Precision, recall, and F1 scores for classification of the relative activity ratios based on zero-shot predictions (i.e., one class defined by 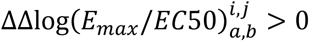, and the other by 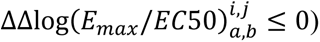 are reported in Figure 8 and in Table S7. The results indicate mediocre performance, with F1 scores of 0.64 (0.59,0.71) for OPRM and 0.49 (0.44,0.60) for D2DR. The low F1 scores are primarily driven by poor recall, which is 0.63 (0.55,0.68) for OPRM and 0.40 (0.31,0.49) for D2DR. Notably, AUCROC cannot be calculated here because class assignment is not based on probabilistic predictions.

### Few-Shot Fine-Tuning Enhances Ligand Bias Prediction For Selected Class A GPCRs

We investigated whether few-shot fine-tuning could enhance the performance of classifiers of relative activity ratio predictions. After fine-tuning the final layer of the classifier separately on 10% of the training data for OPRM and D2DR, we observed a significant improvement in recall (see Figure 8, dark boxes, and Table S7). Specifically, recall for OPRM increased from 0.63 to 0.76 (0.72,0.82) and for D2DR, it improved from 0.40 to 0.70 (0.65,0.79). The improved recall translated into increased F1 scores of 0.84 (0.76,0.94) for OPRM and 0.64 (0.48,0.74) for D2DR, up from 0.64 and 0.49, respectively.

**Figure 8.**
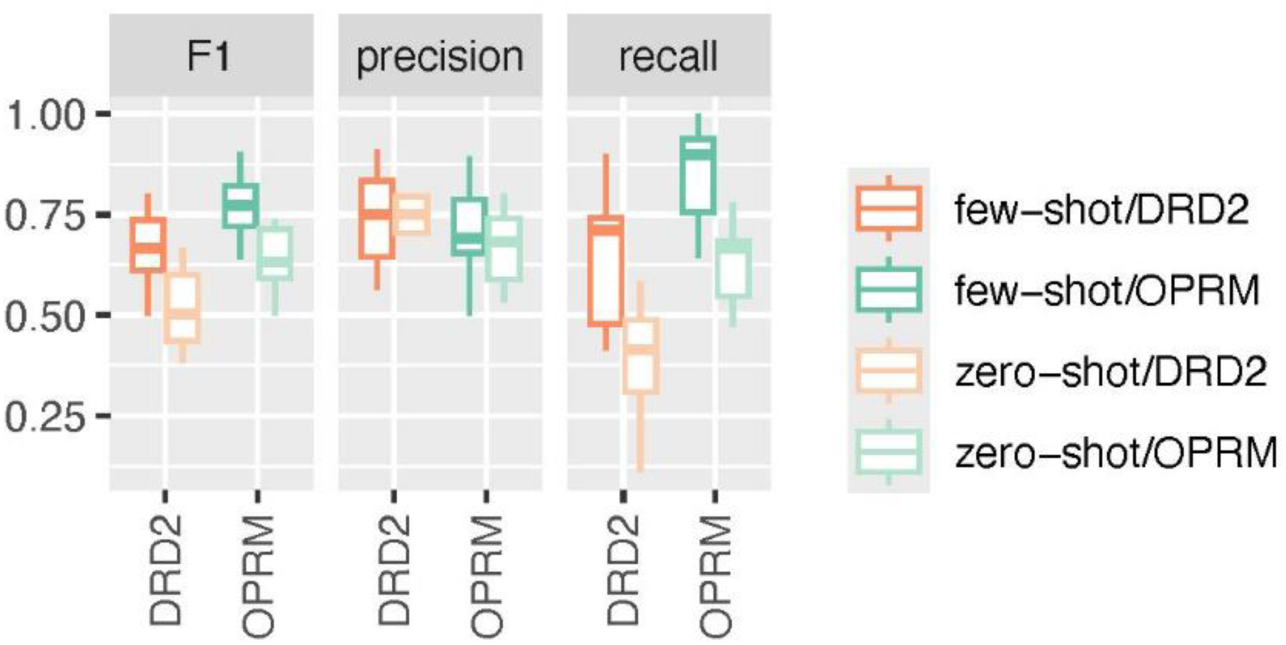
Performance of ligand bias classifiers for D2DR (orange) and OPRM (green), relative to dopamine and DAMGO, respectively. Results for the zero-shot classifier and the model optimized with few-shot fine-tuning are depicted in light and dark hues, respectively.

Given that these classifiers rely on specific reference ligands, we also examined how few-shot fine-tuning impacted the underlying regression models. Figure 9 and Table S8 report data demonstrating significant improvements in the regression performance of models predicting activity ratios of OPRM and D2DR ligands in the G protein and arrestin pathways. These improvements included higher CI, better Pearson and Spearman correlation metrics, and reduced MAE and RMSE. These results suggest that the enhancement of regression models for activity ratio predictions through few-shot fine-tuning could improve model accuracy, as demonstrated here using DAMGO and dopamine as reference ligands. This capability opens the door for more detailed comparisons between multiple reference ligands, further advancing biased agonism research.

**Figure 9.**
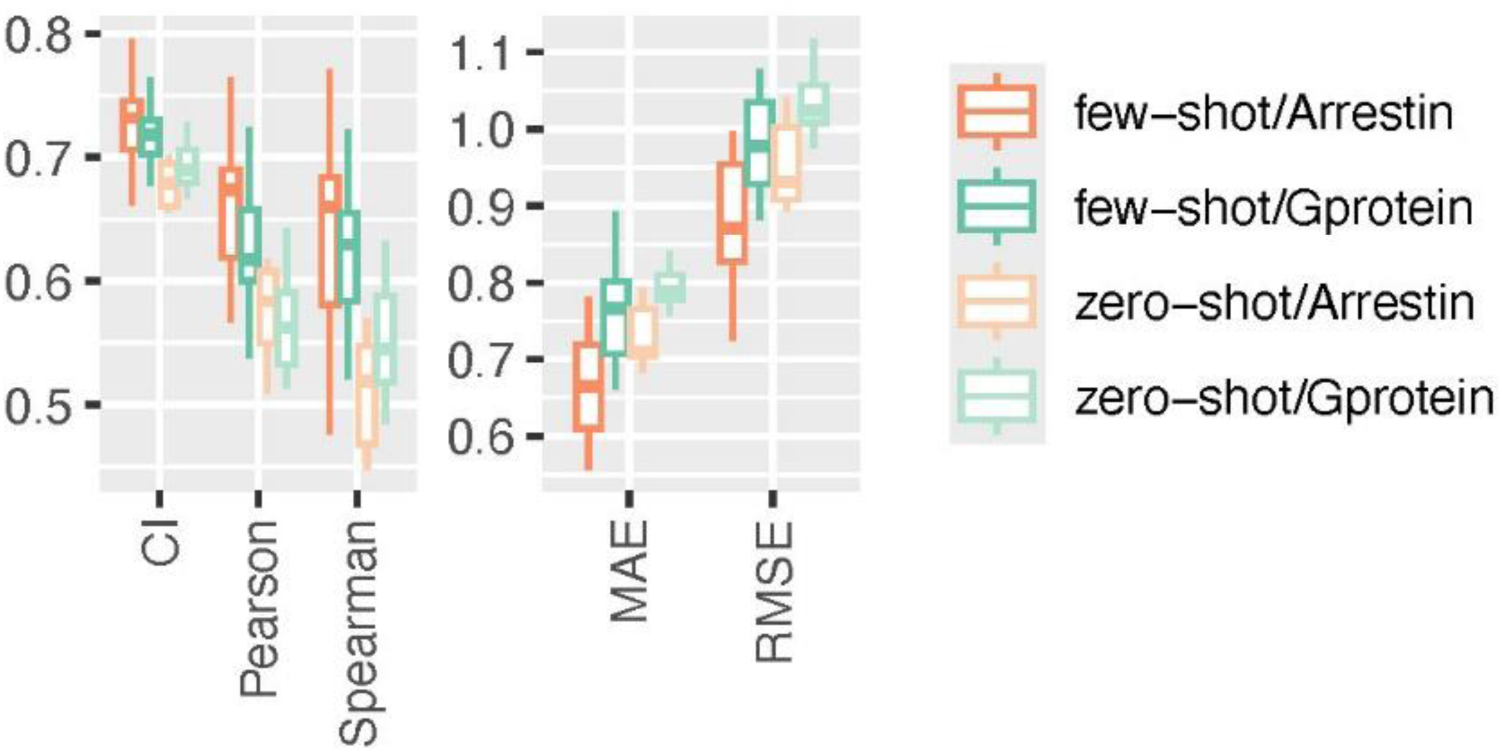
Regression performance for activity ratio predictions of OPRM and DRD2 ligands. The zero-shot (soft hues) and few-shot (bold hues) trained models are compared for predictions in the arrestin (orange) and G protein (green) pathways.

### Deployment of Models for Large Screening

Our fine-tuned transfer learning models, which integrate trained ligand embeddings and fixed embeddings of specific class A GPCR targets, enable the calculation of ligand fingerprints and classification in a massively parallelized manner with minimal computational overhead, thereby facilitating the *in silico* screening of large databases. Fingerprinting a single ligand from the potency-based classifier training set took an average of 8.6 ms on a single core of an x86_64 Intel Xeon W-2255 CPU running at 3.70GHz. This performance extrapolates to ∼14 min for processing one million ligands using a 10-core CPU. Compounds from a large dataset such as the Enamine’s REAL HAC 11-21 subset (*43*), could be fingerprinted with comparable efficacy, averaging 8.4 ms per ligand on the same hardware. Using a single Nvidia RTX A4000 GPU, our specialized classifiers can assign ligand labels at an average speed of 0.012 ms per ligand. This translates to ∼3.3 hours to process one billion ligands, compared to 0.98 ms per ligand when the model needs to also compute the embedding of the target sequence. These benchmarks highlight the efficiency of the presented models in screening large ligand databases, establishing them as effective tools for high-precision prescreening and ligand prioritization across various class A GPCRs. This capability paves the way for the discovery of compounds with potentially improved safety profiles.

## CONCLUSIONS

Transfer learning once again proves to be a powerful strategy for enhancing bioactivity predictions in class A GPCRs. Leveraging the larger dataset of ligand-GPCR activities available for the entire family, we achieved robust precision and recall in predicting active ligands across various data splits. Moreover, a fine-tuned efficacy-based classifier successfully distinguished partial from full agonists, demonstrating superior performance compared to models trained directly on limited efficacy data. However, the performance of this fine-tuned efficacy-based classifier declined when applied to unseen targets. Few-shot fine-tuning on specific receptors significantly improved the performance of efficacy-based classifiers, particularly for GPCR subfamilies with smaller or lower-quality training datasets, underscoring its value in data-limited scenarios.

Predicting ligand activity ratios to assess ligand bias in class A GPCRs remains challenging due to limited data and assay variability, which necessitate relative activity comparisons rather than direct classifications. While initial regression models based on potency data achieved moderate accuracy, their classification performance for ligand bias was suboptimal. However, few-shot fine-tuning significantly improved recall and F1 scores for ligand bias predictions of OPRM and D2DR ligands. Additionally, it enhanced regression metrics for predicting activity ratio values of OPRM and D2DR ligand, using DAMGO and dopamine as reference ligands, respectively. These improvements demonstrate the potential of this approach to deliver more accurate and detailed ligand bias predictions across multiple reference ligands.

Further studies are needed to assess the impact of averaging across different assay types, conditions, and biological endpoints within the G protein- and arrestin-mediated signaling pathways. Despite the noted challenges, the protein language models presented here demonstrate significant potential to accelerate drug discovery for class A GPCRs. By enabling precise predictions of ligand efficacy and bias, as well as supporting efficient large-scale *in silico* screening, these models can streamline compound prioritization and advance the development of novel therapeutics. These models are made available to the scientific community upon request.

## Author Contributions

M.F. conceived, supervised, and funded the research work. D.P. jointly designed the strategy with M.F., wrote scripts, and conducted all computations. M.F. and D.P. analyzed the data and wrote the manuscript.

## Notes

The authors declare no competing financial interest.

## ACKNOWLEDGMENTS

The authors would like to thank Drs. Jonathan Javitch, Robert Lane, and Meritxell Canals for helpful discussions. Computations were supported in part through the computational resources and staff expertise provided by Scientific Computing at the Icahn School of Medicine at Mount Sinai, the Clinical and Translational Science Awards (CTSA) grant UL1TR004419 from the National Center for Advancing Translational Sciences, and the Office of Research Infrastructure of the National Institutes of Health under award number S10OD026880. The content is solely the responsibility of the authors and does not necessarily represent the official views of the National Institutes of Health.

