## Supplemental Tables for "Fine-Tuned Deep Transfer Learning Models for Large Screenings of Safer Drugs Targeting Class A GPCRs"

Table S1……………. pg. 2

Table S2……………. pg. 3

Table S3……………. pg. 4

Table S4……………. pg. 5

Table S5……………. pg. 6

Table S6……………. pg. 7

Table S7……………. pg. 8

Table S8……………. pg. 9

**Table S1.** Dataset sizes for the potency-based and efficacy-based classifiers, as well as the activity ratio regressor. For the classifiers, the sizes of the active and inactive strata are reported, while for the activity ratio dataset, the strata corresponding to the arrestin and G protein effector pathways are provided.

| **Dataset** | **Stratum** | **Datapoints** | **Unique ligands** | **Unique Targets** |
|---|---|---|---|---|
| Potency-based classifier | 0 (inactive) | 168,822 | 74,953 | 531 |
| | 1 (active) | 214,816 | 138,232 | 566 |
| | **Total** | **383,638** | **180,340** | **615** |
| Efficacy-based classifier | 0 (full) | 6,278 | 5,057 | 176 |
| | 1 (partial) | 4,193 | 3,467 | 150 |
| | **Total** | **10,471** | **7,749** | **198** |
| Activity-ratio regressor | Arrestin | 2,304 | 864 | 56 |
| | G Protein | 3,781 | 992 | 62 |
| | **Total** | **6,085** | **1,030** | **64** |

**Table S2.** Size of the potency-based classifier, efficacy-based classifier, and activity ratio regressor datasets for the class A GPCR families with the largest datasets. The total numbers of data points, unique ligands, and unique receptors in each stratum are separated by slashes.

| Family Name | Potency-based classifier | Efficacy-based classifier | Activity-ratio regressor |
|---|---:|---:|---:|
| 5-Hydroxytryptamine receptors | 46,177/23,915/49 | 1,937/1,433/17 | 272/73/7 |
| Opioid receptors | 37,865/15,609/18 | 2,297/1,521/14 | 2,092/357/9 |
| Dopamine receptors | 33,305/17,430/22 | 723/575/8 | 1,191/202/5 |
| Adenosine receptors | 32,395/14,272/23 | 273/201/8 | 315/79/4 |
| Adrenoceptors | 25,605/9,176/42 | 317/196/15 | 360/84/4 |
| Cannabinoid receptors | 20,783/12,270/6 | 1,340/1,062/6 | 668/90/3 |
| Acetylcholine receptors (muscarinic) | 18,678/8,499/18 | 235/207/8 | 177/29/5 |
| Histamine receptors | 16,686/10,704/16 | 133/102/7 | 102/29/2 |
| Chemokine receptors | 14,232/12,632/49 | 36/34/5 | 45/22/2 |
| Prostanoid receptors | 13,093/9,185/27 | 30/30/3 | 116/51/2 |
| Other | 124,819/75,412/345 | 3,150/2,474/107 | 747/191/21 |

**Table S3.** Performance on the random, ligands, and target validation splits for pre-training on the potency-based classification task.

| Split | Precision | Recall | F1 | AUCROC |
|---|---|---|---|---|
| ligands | 0.76 (0.76, 0.76) | 0.89 (0.88, 0.89) | 0.82 (0.82, 0.82) | 0.86 (0.86, 0.87) |
| random | 0.80 (0.80, 0.80) | 0.88 (0.87, 0.88) | 0.84 (0.83, 0.84) | 0.89 (0.89, 0.89) |
| targets | 0.71 (0.71, 0.73) | 0.80 (0.79, 0.82) | 0.76 (0.74, 0.77) | 0.78 (0.78, 0.80) |

**Table S4.** Performance on the random, ligand, and target validation splits for the fine-tuned efficacy-based classification task, as well as for the classifier trained directly on the efficacy dataset (without transfer learning from the potency classification task).

| Train strategy | Split | Precision | Recall | F1 | AUCROC |
|---|---|---|---|---|---|
| fine-tuned | ligands | 0.71 (0.71, 0.72) | 0.74 (0.73, 0.75) | 0.73 (0.73, 0.73) | 0.86 (0.85, 0.86) |
| fine-tuned | random | 0.70 (0.70, 0.70) | 0.75 (0.72, 0.78) | 0.72 (0.71, 0.74) | 0.85 (0.85, 0.86) |
| fine-tuned | targets | 0.57 (0.53, 0.61) | 0.52 (0.41, 0.61) | 0.54 (0.48, 0.60) | 0.68 (0.65, 0.67) |
| no pretrain | ligands | 0.66 (0.64, 0.68) | 0.69 (0.65, 0.73) | 0.66 (0.64, 0.67) | 0.83 (0.82, 0.83) |
| no pretrain | random | 0.66 (0.65, 0.67) | 0.71 (0.69, 0.72) | 0.67 (0.66, 0.67) | 0.83 (0.83, 0.84) |
| no pretrain | targets | 0.56 (0.54, 0.58) | 0.49 (0.46, 0.48) | 0.52 (0.50, 0.55) | 0.67 (0.65, 0.68) |

**Table S5.** F1 scores and training dataset sizes for potency-based and efficacy-based classifiers across random validation sets, organized by GPCR subfamily.

| **Family Name** | **F1 score Potency-based classifier** | **F1 score Efficacy-based classifier** | **Potency-based Training Dataset** | **Efficacy-based Training Dataset** |
|---|---|---|---|---|
| 5-Hydroxytryptamine receptors | 0.87 (0.86, 0.87) | 0.74 (0.72, 0.75) | 46,177 | 1,937 |
| Acetylcholine receptors (muscarinic) | 0.77 (0.77, 0.77) | 0.56 (0.55, 0.58) | 18,678 | 235 |
| Adenosine receptors | 0.79 (0.78, 0.79) | 0.80 (0.81, 0.82) | 32,395 | 273 |
| Adrenoceptors | 0.81 (0.81, 0.81) | 0.72 (0.68, 0.78) | 25,605 | 317 |
| Angiotensin receptors | 0.90 (0.89, 0.90) | 0.86 (0.80, 0.91) | 6,034 | 38 |
| Apelin receptor | 0.89 (0.88, 0.90) | 0.17 (0.00, 0.17) | 843 | 98 |
| Bombesin receptors | 0.89 (0.89, 0.89) | 0.00 (0.00, 0.00) | 1,400 | 109 |
| Cannabinoid receptors | 0.76 (0.76, 0.77) | 0.61 (0.58, 0.63) | 20,783 | 1,340 |
| Chemokine receptors | 0.86 (0.86, 0.87) | 0.71 (0.53, 0.82) | 14,232 | 36 |
| Cholecystokinin receptors | 0.73 (0.73, 0.73) | 0.97 (1.00, 1.00) | 7,460 | 16 |
| Class A Orphans | 0.86 (0.86, 0.87) | 0.88 (0.87, 0.91) | 7,818 | 755 |
| Complement peptide receptors | 0.72 (0.70, 0.73) | 0.96 (1.00, 1.00) | 742 | 10 |
| Dopamine receptors | 0.83 (0.83, 0.83) | 0.77 (0.76, 0.77) | 33,305 | 723 |
| Free fatty acid receptors | 0.79 (0.79, 0.80) | 0.39 (0.36, 0.43) | 3,244 | 354 |
| Ghrelin receptor | 0.88 (0.88, 0.88) | 0.34 (0.00, 0.57) | 3,378 | 123 |
| Histamine receptors | 0.85 (0.85, 0.86) | 0.74 (0.67, 0.82) | 16,686 | 133 |
| Hydroxycarboxylic acid receptors | 0.73 (0.72, 0.74) | 0.00 (0.00, 0.00) | 1,162 | 22 |
| Leukotriene receptors | 0.86 (0.85, 0.86) | 0.89 (0.86, 1.00) | 2,951 | 28 |
| Lysophospholipid (LPA) receptors | 0.76 (0.75, 0.77) | 0.20 (0.00, 0.50) | 833 | 38 |
| Lysophospholipid (S1P) receptors | 0.81 (0.80, 0.82) | 0.40 (0.35, 0.46) | 6,837 | 174 |
| Melanocortin receptors | 0.82 (0.81, 0.82) | 0.23 (0.20, 0.33) | 11,308 | 339 |
| Melatonin receptors | 0.91 (0.90, 0.91) | 0.75 (0.72, 0.78) | 2,244 | 225 |
| Neuropeptide FF/AF receptors | 0.88 (0.85, 0.91) | 0.13 (0.00, 0.00) | 350 | 70 |
| Neuropeptide W/B receptors | 0.55 (0.53, 0.56) | 0.92 (0.91, 1.00) | 473 | 28 |
| Neuropeptide Y receptors | 0.80 (0.80, 0.80) | 0.00 (0.00, 0.00) | 5,661 | 16 |
| Neurotensin receptors | 0.82 (0.82, 0.84) | 0.58 (0.40, 0.67) | 1,235 | 60 |
| Opioid receptors | 0.85 (0.85, 0.85) | 0.73 (0.73, 0.74) | 37,865 | 2,297 |
| Orexin receptors | 0.82 (0.81, 0.82) | 0.00 (0.00, 0.00) | 8,062 | 100 |
| P2Y receptors | 0.78 (0.78, 0.78) | 0.98 (1.00, 1.00) | 3,811 | 33 |
| Prostanoid receptors | 0.82 (0.82, 0.82) | 0.97 (1.00, 1.00) | 13,093 | 30 |
| Relaxin family peptide receptors | 0.82 (0.76, 0.89) | 0.00 (0.00, 0.00) | 337 | 78 |
| Tachykinin receptors | 0.91 (0.90, 0.91) | 0.87 (0.67, 1.00) | 6,320 | 16 |
| Trace amine receptor | 0.95 (0.95, 0.95) | 0.67 (0.59, 0.71) | 3,151 | 139 |
| Urotensin receptor | 0.86 (0.86, 0.87) | 0.38 (0.40, 0.46) | 889 | 138 |
| Vasopressin and oxytocin receptors | 0.82 (0.81, 0.82) | 0.85 (0.79, 0.87) | 7,775 | 94 |

**Table S6.** Regression performance on validation set across 5 random cross-validation splits for predicting log activity ratios. Regression predictions for the arrestin and G protein pathways are reported as the mean with 25%-75% intervals.

| metric | Arrestin | G protein |
|---|---|---|
| MAE | 0.86 (0.86,0.88) | 0.87 (0.86,0.88) |
| RMSE | 1.12 (1.10,1.13) | 1.13 (1.11,1.15) |
| Pearson | 0.57 (0.53,0.60) | 0.58 (0.57,0.59) |
| Spearman | 0.55 (0.52,0.58) | 0.57 (0.56,0.59) |
| CI | 0.69 (0.68,0.71) | 0.70 (0.70,0.71) |

**Table S7.** Performance of the biased ligand classifiers for D2DR and OPRM, relative to dopamine and DAMGO, respectively. Values for the zero-shot classifier and the model optimized through few-shot fine-tuning are reported as mean with 25%-75% intervals across 5 random validation splits.

| **Target** | **metric** | **few-shot** | **zero-shot** |
|---|---|---|---|
| D2DR | F1 | 0.64 (0.48,0.74) | 0.49 (0.44,0.60) |
| | precision | 0.76 (0.72,0.82) | 0.74 (0.70,0.79) |
| | recall | 0.70 (0.65,0.79) | 0.40 (0.31,0.49) |
| OPRM | F1 | 0.84 (0.76,0.94) | 0.64 (0.59,0.71) |
| | precision | 0.64 (0.48,0.74) | 0.66 (0.59,0.74) |
| | recall | 0.76 (0.72,0.82) | 0.63 (0.55,0.68) |

**Table S8.** Regression performance of models predicting activity ratios of OPRM and D2DR ligands. The results obtained using the zero-shot and few-shot trained models are compared side by side.

| Pathway | Metric | Few-shot | Zero-shot |
|---|---|---|---|
| Arrestin | MAE | 0.67 (0.61,0.72) | 0.73 (0.70,0.77) |
| | RMSE | 0.88 (0.83,0.95) | 0.95 (0.91,1.00) |
| | Pearson | 0.66 (0.62,0.69) | 0.58 (0.55,0.61) |
| | Spearman | 0.64 (0.58,0.68) | 0.51 (0.47,0.55) |
| | CI | 0.73 (0.71,0.75) | 0.68 (0.66,0.69) |
| G protein | MAE | 0.76 (0.71,0.80) | 0.80 (0.78,0.81) |
| | RMSE | 0.98 (0.93,1.03) | 1.04 (1.01,1.06) |
| | Pearson | 0.63 (0.60,0.66) | 0.57 (0.53,0.59) |
| | Spearman | 0.62 (0.58,0.65) | 0.55 (0.52,0.59) |
| | CI | 0.72 (0.70,0.73) | 0.69 (0.68,0.71) |
